# Copper intoxication in group B *Streptococcus* triggers transcriptional activation of the *cop* operon that contributes to enhanced virulence during acute infection

**DOI:** 10.1101/2021.03.25.437115

**Authors:** Matthew J. Sullivan, Kelvin G. K. Goh, Dean Gosling, Lahiru Katupitiya, Glen C. Ulett

## Abstract

Bacteria can utilize Copper (Cu) as a trace element to support cellular processes; however, excess Cu can intoxicate bacteria. Here, we characterize the *cop* operon in group B streptococcus (GBS), and establish its role in evasion of Cu intoxication and the response to Cu stress on virulence. Growth of GBS mutants deficient in either the *copA* Cu exporter, or the *copY* repressor, were severely compromised in Cu-stress conditions. GBS survival of Cu stress reflected a mechanism of CopY de-repression of the CopA efflux system. However, neither mutant was attenuated for intracellular survival in macrophages. Analysis of global transcriptional responses to Cu by RNA-sequencing revealed a stress signature encompassing homeostasis of multiple metals. Genes induced by Cu stress included putative metal transporters for manganese import, whereas a system for iron export was repressed. In addition, *copA* promoted the ability of GBS to colonize the blood, liver and spleen of mice following disseminated infection. Together, these findings show that GBS *copA* mediates resistance to Cu intoxication, via regulation by the Cu-sensing transcriptional repressor, *copY*. Cu stress responses in GBS reflect a transcriptional signature that heightens virulence and represents an important part of the bacteria’s ability to survive in different environments.

**Importance:** Understanding how bacteria manage cellular levels of metal ions, such as copper, helps to explain how microbial cells can survive in different stressful environments. We show how the opportunistic pathogen group B Streptococcus (GBS) achieves homeostasis of intracellular copper through the activities of the genes that comprise the cop operon, and describe how this helps GBS survive in stressful environments, including in the mammalian host during systemic disseminated infection.

## Introduction

Copper (Cu) is the most reactive of the biologically-relevant first-row *d-*block transition metals. It can readily displace other metals, including Zinc (Zn), Nickel (Ni), Cobalt (Co), Iron (Fe) and Manganese (Mn) from metalloproteins in which these are bound (1, 2). In cells across all Kingdoms of life, Cu-dependent enzymes are pivotal to many essential processes that underpin physiologic biochemical reactions, owing to its ability to redox cycle between Cu(I) and Cu(II) states (3). When present in excess, however, Cu is hazardous to cellular processes and macromolecules due to effects of localized free-radical damage (4). A key role for Cu in bacterial cell biology encompasses the host-pathogen interface where a balance between Cu usage and avoidance of Cu intoxication resulting from host antimicrobial responses must be attained for bacteria to survive, as reviewed elsewhere (5). At this interface, human immune cells can mobilize Cu in a defence response to infection that culminates in exposure of intracellular bacteria to antimicrobial levels of Cu that kill the invading pathogen (6-9). Thus, bacterial responses to Cu stress and mechanisms of resistance to metal-ion intoxication have emerged as important facets of bacterial disease pathogenesis (10).

There are several mechanisms that enable bacteria to tolerate excess extracellular Cu, and these have been characterized in several pathogenic species. These mechanisms, which represent survival strategies against Cu intoxication (10) typically involve Cu-transporting P-type ATPases, exemplified by CopA, which detoxify Cu by exporting it from the bacterial cytosol (11-13). This mechanism has been described in gram-negative and gram-positive bacteria, and among *Streptococcaceae* (14-16), including pneumococci and *Streptococcus pyogenes* (17).

An opportunistic streptococcal pathogen of humans and animals for which resistance to excess extracellular Cu has not been described is *Streptococcus agalactiae*, also known as group B streptococcus (GBS). This organism, unlike other *Streptococcaceae*, is associated with a comparatively broad host-range that encompasses humans, cattle and fish (18). In humans, GBS is a major cause of invasive infection in infants <3 months of age (19), and causes a more diverse range of disease aetiologies compared to other streptococci, including meningitis, pneumonia, skin and soft-tissue infections, sepsis, arthritis, osteomyelitis, urinary tract infection, and endocarditis (19). GBS has several virulence factors that enable survival in cytotoxic environments, such as acid stress, oxidative stress, and during host colonization, as reviewed elsewhere (20). Cellular tolerance to Cu stress and mechanisms of responding to and surviving Cu intoxication have not been defined in GBS.

In this study, we characterise a system that enables Cu efflux in GBS, encompassing a *copA*-homologue, which we show mediates control of Cu efflux in the bacteria. We show this system affects survival, growth and virulence of GBS in the mammalian host.

## Results

### Transcriptomic profiling of GBS responses to extracellular Cu

To dissect the global response of GBS to extracellular Cu stress we performed RNA sequencing (RNASeq) to define the complete primary transcriptional response to Cu. RNASeq analysis of mid-log phase cells of GBS strain 874391 grown in the presence of 0.5 mM Cu (a sub-inhibitory level; see below) for 2.5h (compared to control cultures that were grown without supplemental Cu) revealed a surprisingly modest Cu-responsive transcriptome. The response comprised 18 transcriptional responses, defined as significant based on -2≥ FC ≥2 (FDR < 0.05, *n*=4 biological replicates), of 11 up-regulated transcripts and 7 down-regulated transcripts (Figure 1, Table 1). The most significantly up-regulated were homologues of the *copY-copA-copZ* gene cluster (3.6-4.0-fold), which encode the putative CopY (Cu-binding transcriptional repressor), CopA (P-type ATPase Cu-efflux system) and CopZ (Cu-chaperone) proteins. To validate the expression levels of selected targets identified in RNASeq, we targeted *pcl1*, *hvgA* and *copY* in qRT-PCR assays, confirming near-identical fold-change values to RNASeq analyses (Figure 1B). The products of *copY*, *copA* and *copZ* in GBS (Figure 2A) are moderately conserved compared to other Lactobacillales, including *S. mutans, S. thermophilus, S. pyogenes, S. pneumoniae*, and *Enterococcus*(Figure 2B, ranked by CopA similarity) for which a homologous Cu management system is defined (9, 13-15). Together, these findings provide transcriptional and comparative insights that support a proposed model of Cu efflux in GBS (Figure 2C).

**FIG 1.**
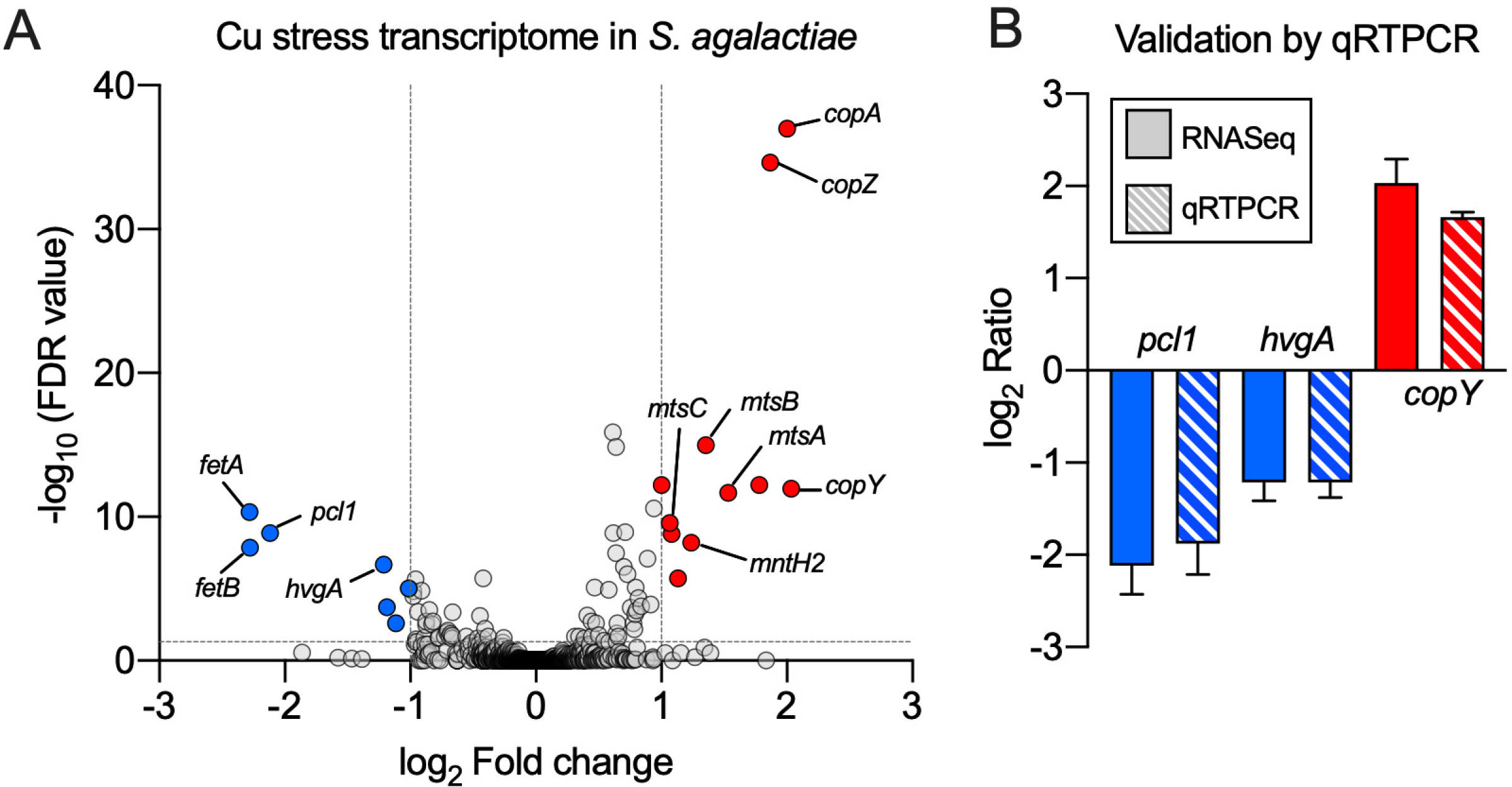
Global transcriptomic analysis of GBS in response to Cu stress. Volcano plot showing data from RNASeq analysis of WT GBS cultures exposed to 0.5 mM Cu compared to non-exposed controls. Transcripts detected as up- or down-regulated in response to Cu (*n*=4, >± 2-fold, FDR <0.05) are highlighted in red and blue, respectively. Dotted lines show False discovery rate (FDR; q-value) and fold-change cut-offs. Grey points indicate genes that were not significant changed in expression, according to these analysis cut-offs. Selected genes are identified with black lines. B, Validation of RNASeq data. Expression ratio (Fold-change) of *pcl1*, *hvgA* and *copY* quantified by qRTPCR in THB medium containing 0.5 mM Cu compared to THB without Cu. Ratios for qRTPCR were normalized using housekeeping *dnaN* and RNASeq ratios were calculated using DESeq2. Bars show means and S.E.M from 4 independent experiments.

**FIG 2.**
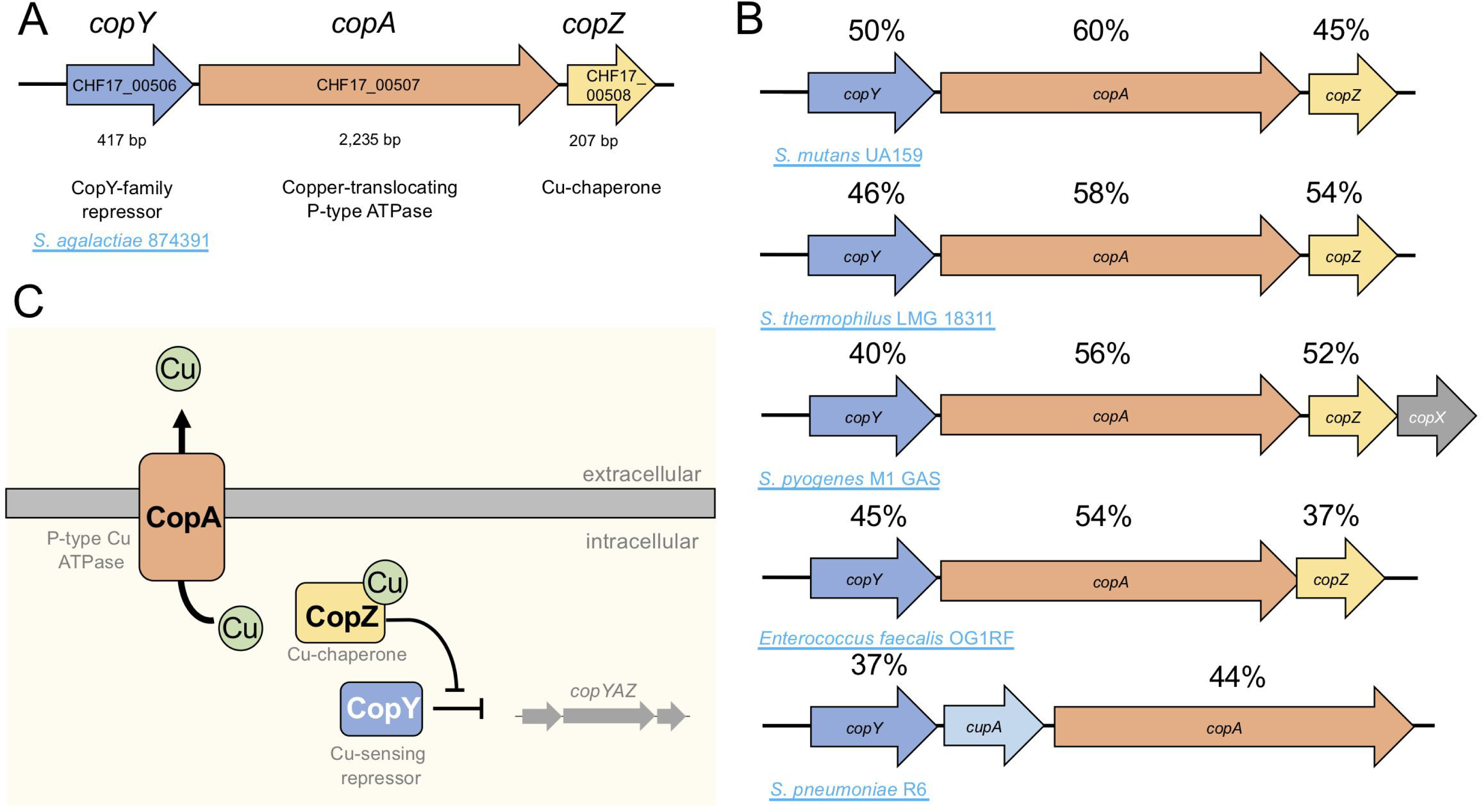
Organisation of the *copY-copA-copZ* locus in GBS. A, *copY-copA-copZ* are adjacent in the GBS genome and likely controlled by the promoter-proximal *copY,* encoding a putative Cu-sensing repressor. Locus tags from the GBS 874391 genome are indicated. B, Distribution of homologous *cop* genes in other Lactobacillales, arranged by percentage identity of amino acid sequence to CopA of *S. agalactiae*. C, Model of Cu efflux in GBS highlighting CopA as transmembrane Cu exporter, transcriptionally repressed by CopY in the absence of Cu, which is likely inhibited by the Cu-binding chaperone protein CopZ. Figure based on previous studies in other lactobacillales (53, 63).

**TABLE 1.** Transcriptional signature of Cu intoxication in GBS

| **Locus_**<br>**tag<sup>†</sup>** | **Genbank**<br>**accession** | **Label** | **Annotation** | **FC** | **FDR** |
| --- | --- | --- | --- | --- | --- |
| 02565 | ASZ00809.1 | *copY* | CopY/TcrY family copper transport repressor | 4.1 | 1.09E-12 |
| 02575 | ASZ00811.1 | *copZ* | carbonate dehydratase | 4.0 | 1.02E-37 |
| 02570 | ASZ00810.1 | *copA* | copper-translocating P-type ATPase | 3.7 | 2.25E-35 |
| 10970 | ASZ02392.1 |  | peptidoglycan-binding protein LysM | 3.4 | 6.13E-13 |
| 07965 | ASZ01821.1 | *mtsA* | metal ABC transporter substrate-binding protein | 2.9 | 2.15E-12 |
| 07960 | ASZ01820.1 | *mtsB* | metal ABC transporter ATP-binding protein | 2.6 | 1.02E-15 |
| 10190 | ASZ02237.1 | *mntH2* | divalent metal cation transporter | 2.4 | 6.34E-09 |
| 10965 | ASZ02391.1 |  | transglycosylase | 2.2 | 1.87E-06 |
| 09125 | ASZ02032.1 |  | CHAP domain-containing protein | 2.1 | 1.58E-09 |
| 07955 | ASZ01819.1 | *mtsC* | metal ABC transporter permease | 2.1 | 2.62E-10 |
| 10305 | ASZ02261.1 |  | PAP2 family protein | 2.0 | 6.13E-13 |
| 08045 | ASZ01834.1 |  | glycosyltransferase family 2 protein | -2.0 | 9.51E-06 |
| 08200 | ASZ01861.1 |  | branched-chain amino acid ABC transporter substrate-binding protein | -2.3 | 1.98E-04 |
| 10430 | ASZ02284.1 | *hvgA* | pathogenicity protein | -2.3 | 2.18E-07 |
| 04720 | ASZ01199.1 | *pcl1* | membrane protein | -4.3 | 1.38E-09 |
| 04185 | ASZ01095.1 | *fetB* | iron export ABC transporter permease subunit, FetB | -4.9 | 1.39E-08 |
| 04180 | ASZ01094.1 | *fetA* | ABC transporter ATP-binding protein FetA | -4.9 | 4.58E-11 |

<sup>†</sup> denotes locus tag of *S. agalactiae* 874391, preceded by CHF17\_RS; FC, Fold-
Change; FDR, False Discovery Rate

In addition to *copYAZ*, GBS up-regulated putative metal transporters for Mn import (*mtsABC*, *mntH2*), and down-regulated an iron (Fe) export system *(fetAB;* Figure 1, Table 1). In addition, we detected significant down-regulation of *pcl1*, that encodes a putative membrane protein, and the gene for hypervirulence-associated factor, *hvgA*.

### Roles of copA and copY in Cu-resistance in GBS

To functionally characterize two major elements of the Cu-responsive transcriptome in GBS, we targeted *copA* and *copY* by generating isogenic deletion mutants and comparing growth of these with the WT. In a nutrient-rich medium (Todd Hewitt Broth; THB) supplemented with increasing amounts of Cu, we observed no growth inhibitory effects of high Cu (up to 1.5 mM) for WT GBS; the lag phases, growth rates, and final biomass yields were equivalent for the WT comparing different levels of Cu (Figure 3). In contrast, growth of Δ*copA* GBS was severely attenuated by Cu stress ≥ 1 mM; complementation of the *copA* mutation restored the phenotype to WT (Figure 3). Deletion of *copY*, encoding a putative Cu-dependent repressor of *copA*, had no effect on the growth rate or lag phase in THB supplemented with Cu but significantly affected final biomass yield of cultures (*D* at 600nm at 18h; Supplementary Figure S1). Together, this shows that *copA* confers cellular resistance to Cu stress in GBS in a manner that supports bacterial growth in nutritive conditions.

**FIG 3.**
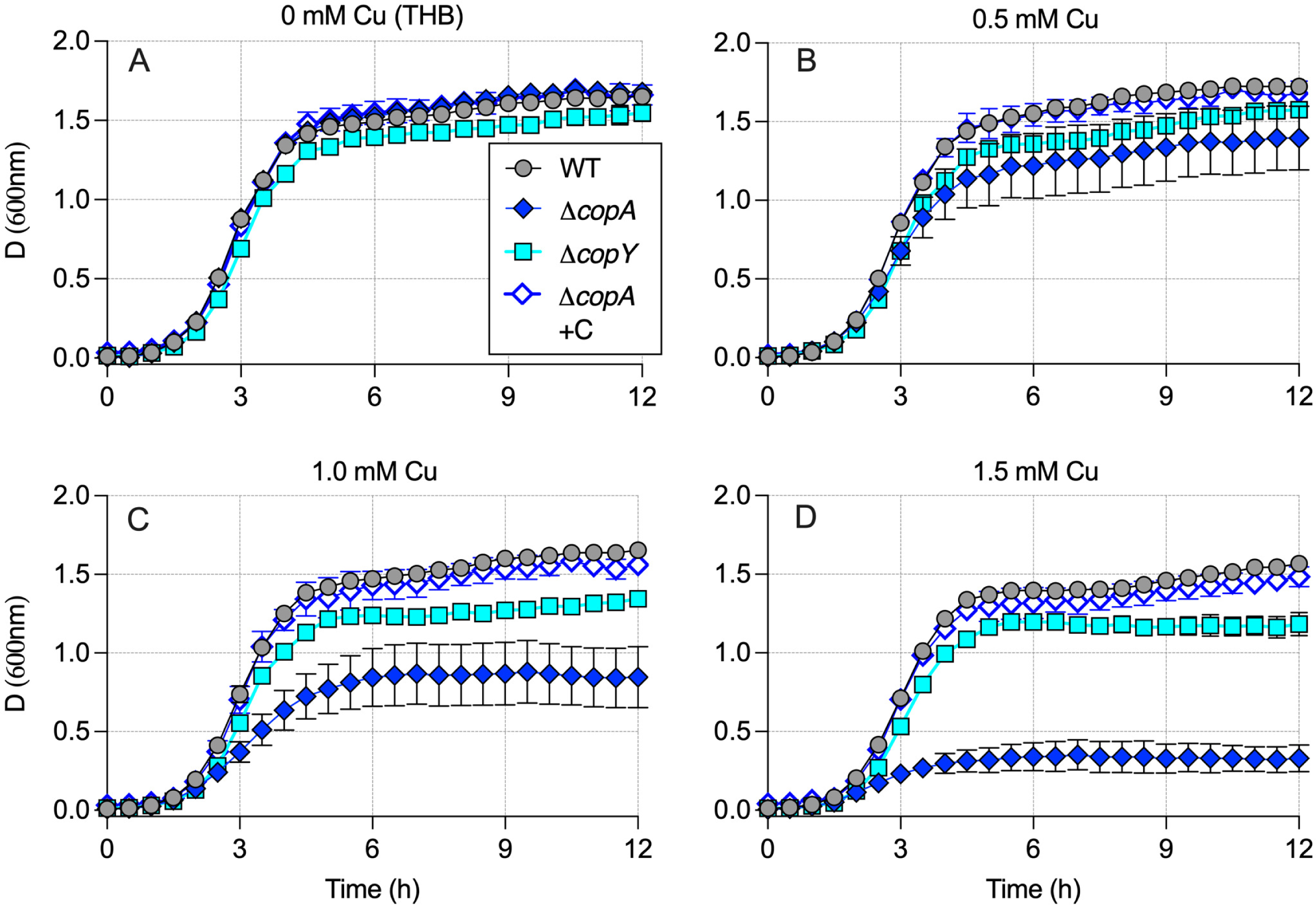
Growth curve analyses of GBS cultured in nutrient-rich THB medium (A), or in THB supplemented with 0.5 mM Cu (B), 1.0 mM Cu (C) or 1.5 mM Cu (D), comparing WT, Δ*copA* or Δ*copY* strains. Complementation of *copA* mutation (Δ*copA*+C) restored growth of GBS to WT levels in conditions of Cu stress. Points and bars show mean and S.E.M of several independent experiments (5 for WT, Δ*copA* and Δ*copY*, 2 for complemented strain) monitoring attenuance at 600nm.

### Temporal- and Concentration-Dependent Bactericidal effects of Cu towards GBS

We examined the effects of Cu towards survival of GBS in nutrient limited conditions by using a minimal Chemically-Defined Medium (CDM), in consideration of prior studies of antibacterial activity in minimal media (21, 22), the influence of culture media (23, 24), and possible cell-protective effects from buffering agents, *e.g.* glutathione (15). We observed that the level of Cu required to inhibit GBS growth in CDM was markedly lower than THB (Supplementary Figure S2); for example, the growth of Δ*copA* GBS was significantly attenuated in CDM in the presence of 0.5 mM supplemental Cu, a level that caused only slight inhibition of growth in THB (Figure 3). To more precisely define the bactericidal effects of Cu towards GBS in CDM, we performed time-kill curves using 1-5 x 10^6^ CFU/mL exposed to Cu concentrations ranging between 50 μM and 1 mM. We observed potent killing effects that depended on both Cu concentration and time with ≥ 0.5 mM Cu significantly killing WT GBS after 3h of exposure (Figure 4, WT); after 24h, there was a ∼230-fold reduction in CFU/mL at 0.5 mM Cu (0 mM Cu = 5.6 Log_10_ CFU/mL, 0.5 mM CFU/mL = 3.2 Log_10_ CFU/mL), and no viable GBS remained in cultures exposure to 1 mM Cu. The Δ*copA* strain was significantly more susceptible to Cu-induced toxicity compared to the WT; Cu levels above ≥ 0.5 mM significantly reduced the number of viable Δ*copA* GBS beginning as early as 1h exposure (Figure 4, Δ*copA*); the degree of bacterial killing was significant at the 24h timepoint for ≥ 0.5 mM Cu (Figure 4). Interestingly, at the 6h timepoint, Cu at ≥ 0.5 mM did not affect viability of Δ*copY* GBS, compared to the WT strain (Figure 4, Δ*copY*), a hyper-resistance phenotype that was not apparent at the 24 h timepoint. Together, these findings are consistent with a role for CopA in resisting Cu stress, and CopY as Cu-dependent repressor of *copA*.

**FIG 4.**
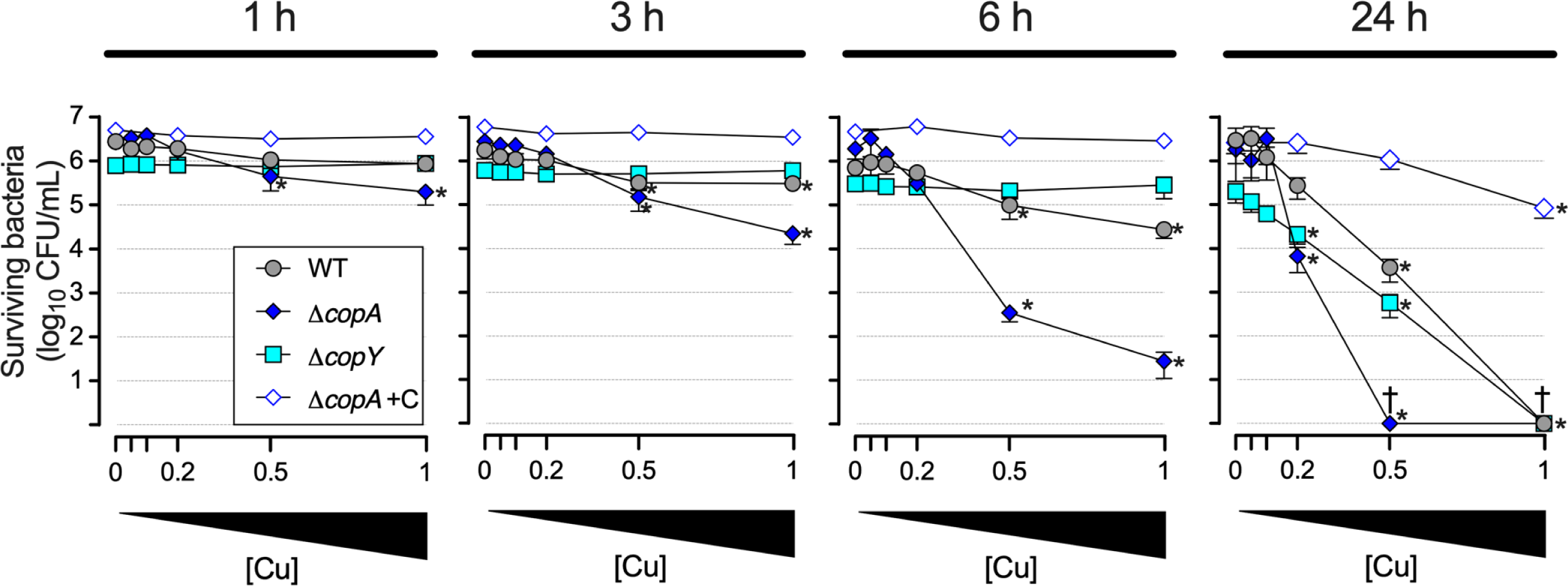
Bactericidal effect of Cu on GBS viability. Time-kill assays comparing WT, Δ*copA* or Δ*copY* GBS (and complemented strain Δ*copA*+C), incubated in CDM or in CDM supplemented with 0.05, 0.1, 0.2, 0.5 or 1 mM Cu. Viable cells were quantified at 1h, 3h, 6h and 24h post incubation. ^†^ Viable Cell counts of 0 CFU/mL were assigned a value of 1 to plot on log_10_ y-axes. Points and bars show mean and S.E.M of 4 independent experiments. Data were analysed by One-way ANOVA with Holm Sidak Multiple comparisons. Significance markers represent comparisons of the number of surviving bacteria in conditions of the defined Cu concentration [Cu] vs the number of bacteria in the non-exposed control (0 mM Cu) for the same time point and same strain (* P < 0.05).

### Regulation of Cu efflux by CopY, and accumulation of metals during Cu stress

We next sought to examine the control of Cu-responses in GBS at the transcriptional level. The capacity of Cu stress to induce expression of *copA* for Cu export was examined by analysing *copA* expression by qRTPCR in GBS exposed to Cu concentrations ranging from 0.25-1.5 mM Cu in THB. GBS significantly upregulated *copA* in response to Cu (3.7-fold – 14.2-fold) in a manner that was titratable with the Cu concentration (Figure 5A). To ascertain the role of CopY as a putative Cu-responsive repressor of *copA* expression in GBS, we quantified *copA* mRNA transcripts in the Δ*copY* mutant exposed to 0.5 mM Cu. This level of Cu was carefully chosen as sub-inhibitory to enable comparisons independent of metabolic state and therefore limiting any potential bias from possible discordant Cu stress between WT and mutants with varied resistance phenotypes. Deletion of *copY* resulted in severe de-regulation of *copA* expression, causing a ∼207±45-fold increase of *copA* transcripts in the *copY* background (Figure 5B). Thus, these data establish that GBS *copY* represses *copA* in the absence of Cu, and are consistent with a model of CopY-mediated regulation of *copA* that is responsive to intracellular accumulation of Cu.

**FIG 5.**
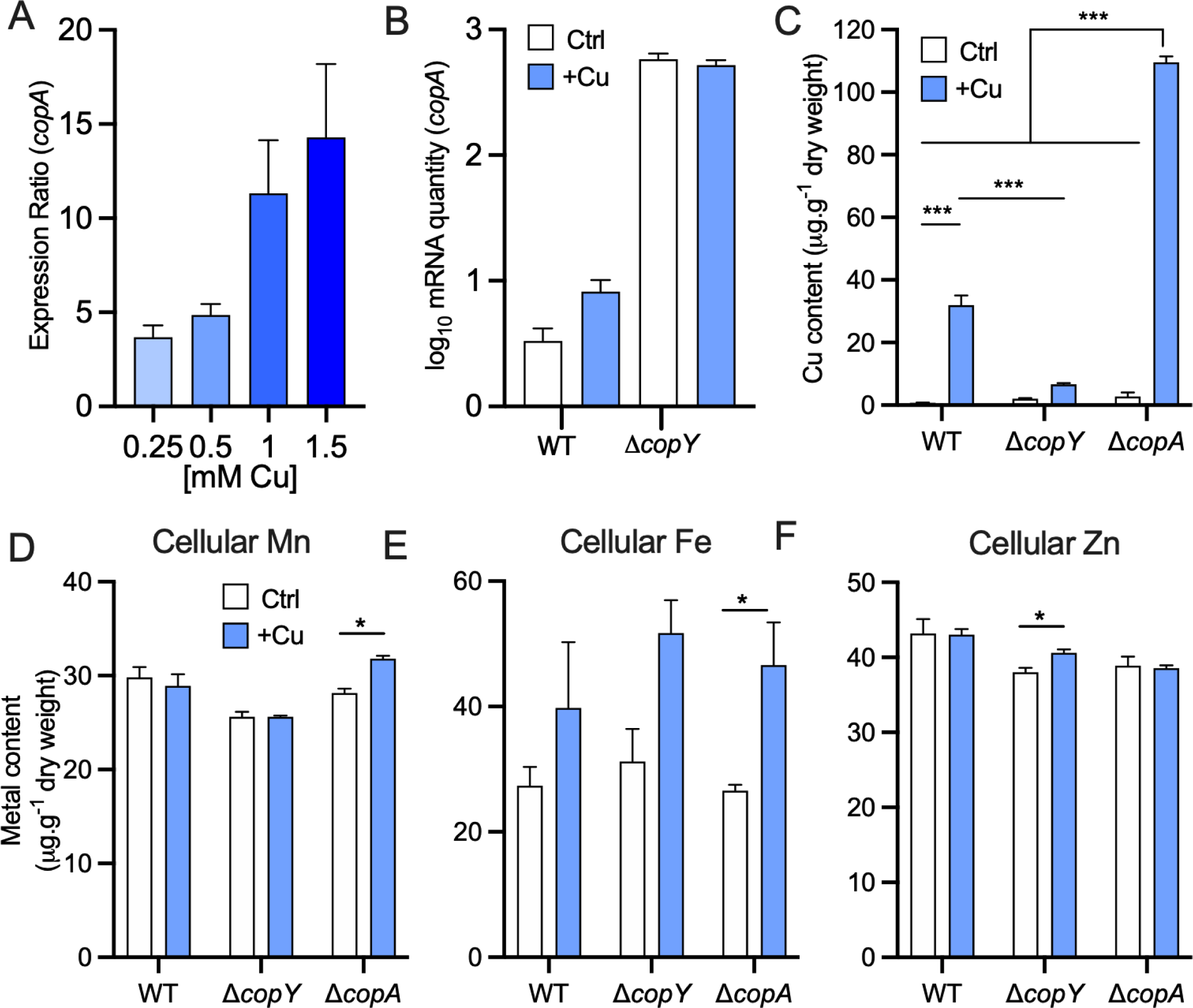
Expression analysis of *copA* and intracellular metal content in GBS strains. A, Expression ratio (Fold-change) of *copA* quantified by qRT-PCR in THB medium containing 0.25, 0.5, 1.0 and 1.5 mM Cu, compared to THB without Cu. B, Relative *copA* transcripts were quantified in WT and Δ*copY* strains with and without Cu supplementation (0.5 mM) to demonstrate de-regulation of *copA* expression in the Δ*copY* background. Intracellular accumulation of Cu (C), Mn (D), Fe (E) and Zn (F) was compared with and without Cu supplementation (0.5 mM) in WT, Δ*copY* and Δ*copA* strains. Ratios in A were calculated as described previously (35) using C_T_ values, primer efficiencies and housekeeping *dnaN*. Bars show means and S.E.M from 3-4 independent experiments and compared by One-way ANOVA with Holm-Sidak multiple comparisons (* P < 0.05; *** P <0.001). Ctrl: Control condition (i.e., no supplemental Cu in media). The ANOVA P value from comparisons of between strains was 0.06 for Cellular Fe and 0.01 for Cellular Zn; subsequent pairwise Student’s *t*-tests were used to compare each strain in the Ctrl and Cu condition (* P < 0.05).

To examine the impact of *copA* and *copY* mutations on accumulation of Cu within the cell we used the equivalent conditions per RNASeq and qPCR assays, exposing GBS to 0.5 mM Cu in THB and measuring total Cu content of cells compared to non-exposed controls (Ctrl). Inductively coupled optical emission spectroscopy (ICP-OES) demonstrated that standard THB contained 0.2±0.08 µM Cu, reflecting trace amounts in the medium. In the absence of supplemental Cu, WT GBS limited intracellular Cu content such that only 0.8±0.1 µg Cu. g dry weight^-1^ were detected in cultures grown in THB. However, exposure of WT GBS to 0.5 mM Cu resulted in a significant increase in intracellular Cu to 31.9±3.1 µg Cu. g dry weight^-1^ (Figure 5C). Strikingly, Δ*copY* GBS exhibited significantly less cellular Cu upon exposure to Cu (6.7±0.4 µg Cu. g dry weight^-1^), consistent with the observation that transcription of the Cu-exporter CopA is significantly elevated in this mutant. In addition, we noted significant accumulation of cellular Cu in the Δ*copA* strain (109.6±1.9 µg Cu. g dry weight^-1^), confirming a Cu-efflux role for *copA*. Interestingly, we noted modest but significant increases in Mn and Fe in the Δ*copA* strain, and Zn in the Δ*copY* strain due to Cu stress (Figure 5D-F).

To extend our findings comparing mutant strains to WT at a level of Cu that is sub-inhibitory to the Δ*copA* strain, we next undertook experiments to investigate the impact of severe Cu stress on WT GBS; by repeating metal accumulation analyses using cells grown in 1.5 mM Cu. We detected modest but statistically significant accumulation of Fe inside GBS cells exposed to 1.5 mM Cu, co-occurring with significantly higher Cu levels; however, no difference in Mn or Zn content (Supplementary Figure S3). Fe status has been reported to influence bacterial resistance to peroxide stress (25) and therefore we also undertook experiments to investigate the consequence of Cu toxicity on resistance to oxidative stress using assays of susceptibility to hydrogen peroxide (H_2_O_2_). These experiments demonstrated a significant reduction in the viability of Δ*copA* GBS exposed to H_2_O_2_, which was dependent on pre-exposure to Cu (Supplementary Figure S4). Thus, GBS cells undergoing Cu stress are rendered significantly more susceptible to oxidative stress.

### Role of CopA in macrophage killing of GBS

To examine whether the CopA Cu efflux system in GBS supports survival of the bacteria in phagocytes we performed antibiotic protection assays with human monocyte-derived macrophage-like cells. Macrophages were infected with WT, Δ*copA* GBS or its complemented strain for 1h, and antibiotics were added to kill extracellular bacteria. Viable intracellular GBS were quantified at 1h post-antibiotic addition, and at 24h and 48h to assess intracellular survival. We performed assays comparing culture conditions of standard RPMI medium to medium that was supplemented with 20 μM Cu to ensure adequate availability of Cu for host cellular responses. These assays demonstrated significant reductions in the numbers of viable GBS over the time course (24 to 48h); however, we did not detect any significant differences between the numbers of WT and Δ*copA* GBS recovered from the host cells at any time point (Figure 6). The numbers of bacteria recovered comparing the WT and Δ*copA* mutant were equivalent regardless of the presence of supplemental Cu in the medium. Thus, under these conditions, *copA* does not contribute to the intracellular survival of GBS in human macrophage-like cells.

**FIG 6.**
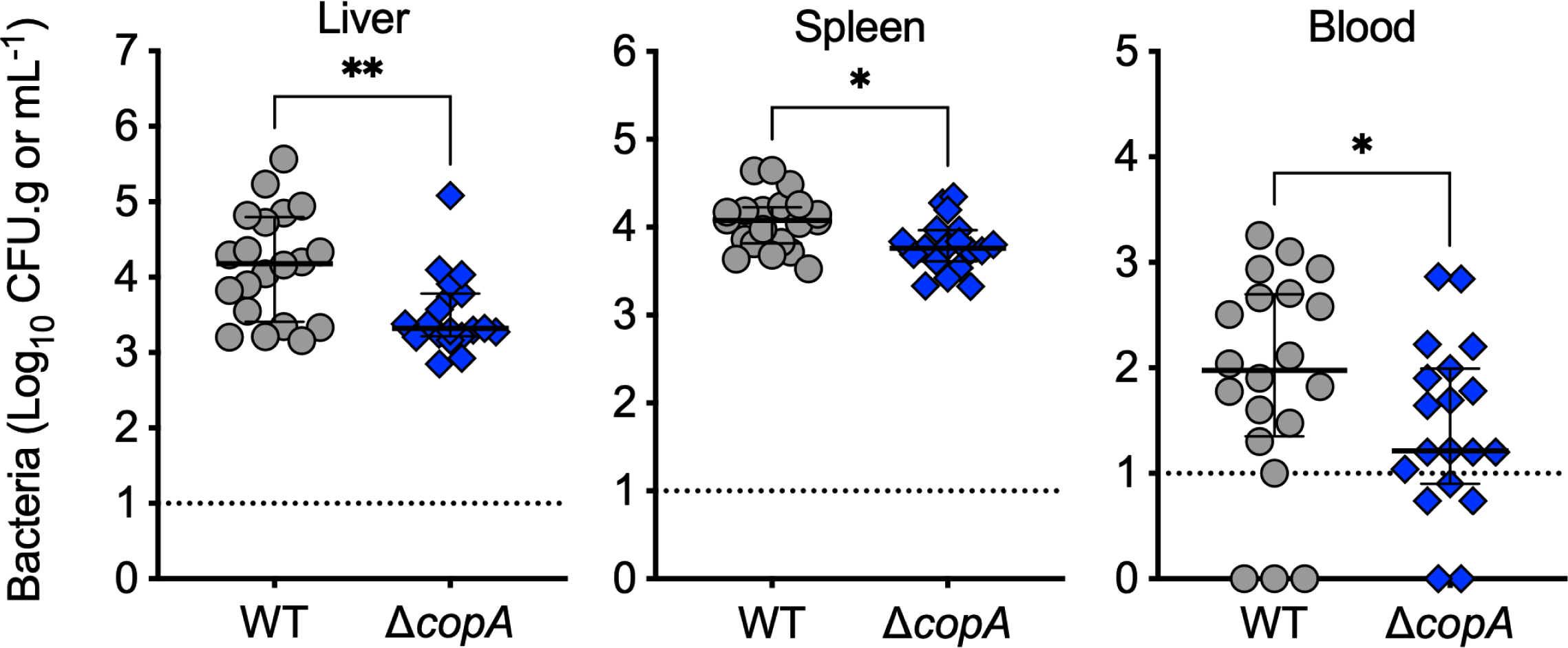
Interactions of GBS with human macrophage-like cells. Gentamicin protection assays with WT and Δ*copA* strains in human (U937 monocyte-derived macrophage-like) macrophages in media with and without 20 μM supplemental Cu. Surviving bacteria are expressed as CFU/mL, indicating the number of intracellular bacteria at 1h after antibiotic treatment, and at 24h and 48h post infection (h.p.i). Data are means and S.E.M of 3-4 independent experiments.

### GBS CopA contributes to virulence in vivo

To examine the contribution of Cu efflux to GBS virulence, we used a murine model of disseminated infection (26). In mice challenged with 10^7^ GBS, we detected significantly fewer Δ*copA* mutant in the liver (median of 3.5 vs 4.2 log_10_ CFU.g tissue^-1^; P= 0.005), spleen (median of 3.8 vs 4.1 log_10_ CFU.g tissue^-1^ P= 0.013) and blood (median of 1.4 vs 1.9 log_10_ CFU.mL^-1^; P= 0.044) compared to the WT at 24h post-inoculation (Figure 7). However, no significant differences were observed between the counts of the WT and Δ*copA* mutant in several other tissues, including the brain, heart, lungs, and kidneys (Supplementary Figure S5). These data support a modest but significant role for cellular management of Cu via *copA* in GBS in supporting disseminated infection *in vivo*.

**FIG 7.**
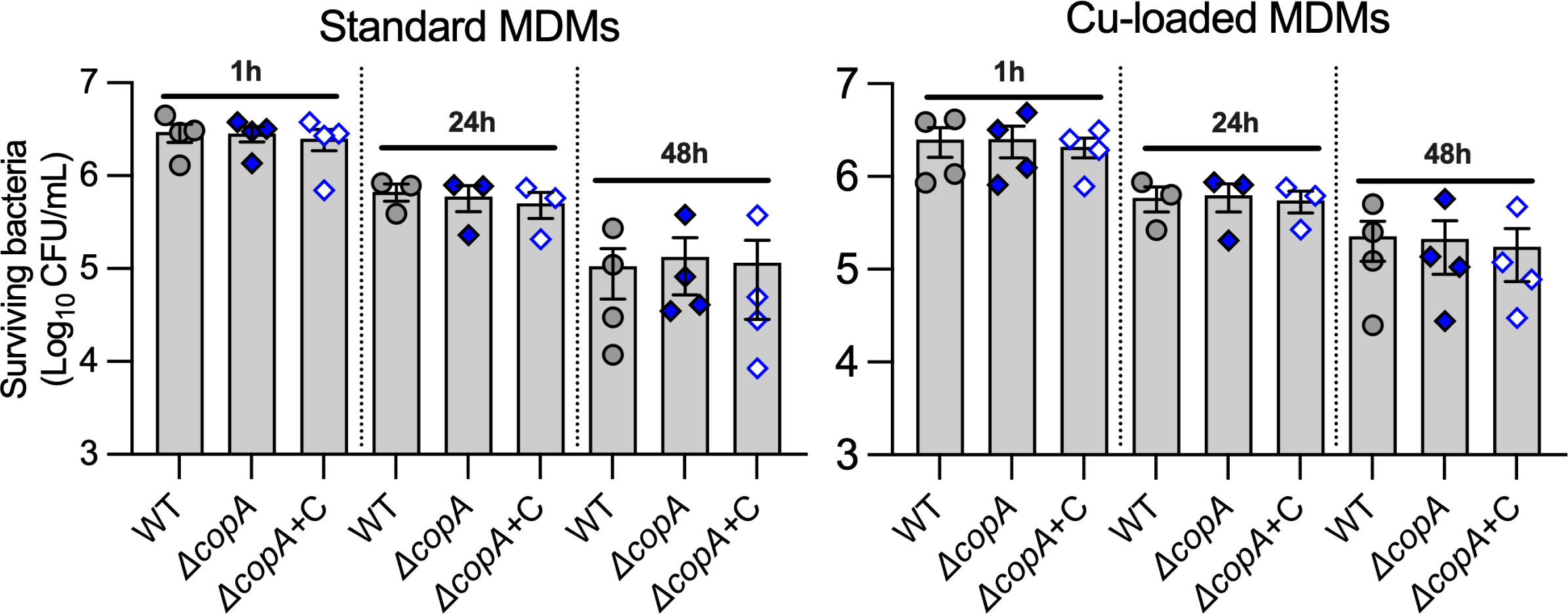
Virulence of WT (grey circles) or Δ*copA* (blue diamonds) GBS in a mouse model of disseminated infection. C57BL/6 mice (6-8 weeks old) were intravenously injected with 10^7^ bacteria; bacteremia and disseminated spread to liver and spleen were monitored at 24h post infection. CFU were enumerated and counts were normalized using tissue mass in g. Viable Cell counts of 0 CFU/mL were assigned a value of 1 to enable visualisation on log_10_ y-axes. Lines and bars show median and interquartile ranges and data are pooled from 2 independent experiments each containing n=10 mice compared using Mann-Whitney U-tests (*P < 0.05, **P < 0.01).

## Materials and Methods

### Bacterial strains, plasmids and growth conditions

GBS, *E. coli* and plasmids used are listed in Table 2. GBS was routinely grown in Todd-Hewitt Broth (THB) or on TH agar (1.5% w/v). *E. coli* was grown in Lysogeny Broth (LB) or on LB agar. Routine retrospective colony counts were performed by plating dilutions of bacteria on tryptone soya agar containing 5% defibrinated horse blood (Thermo Fisher Scientific). Media were supplemented with antibiotics (spectinomycin (Sp) 100μg/mL; chloramphenicol (Cm) 10 μg/mL), as indicated. Growth assays used 200μL culture volumes in 96-well plates (Greiner) sealed using Breathe-Easy® membranes (Sigma-Aldrich) and measured attenuance (*D*, at 600nm) using a ClarioSTAR multimode plate reader (BMG Labtech) in Well Scan mode using a 3mm 5x5 scan matrix with 5 flashes per scan point and path length correction of 5.88mm, with agitation at 300rpm and recordings taken every 30min. Media for growth assays were THB and a modified Chemically-Defined Medium (CDM) (27) (with 1g/L glucose, 0.11g/L pyruvate and 50μg/L L-cysteine), supplemented with Cu (supplied as CuSO_4_) as indicated. For attenuance baseline correction, control wells without bacteria were included for Cu in media alone.

**TABLE 2.**
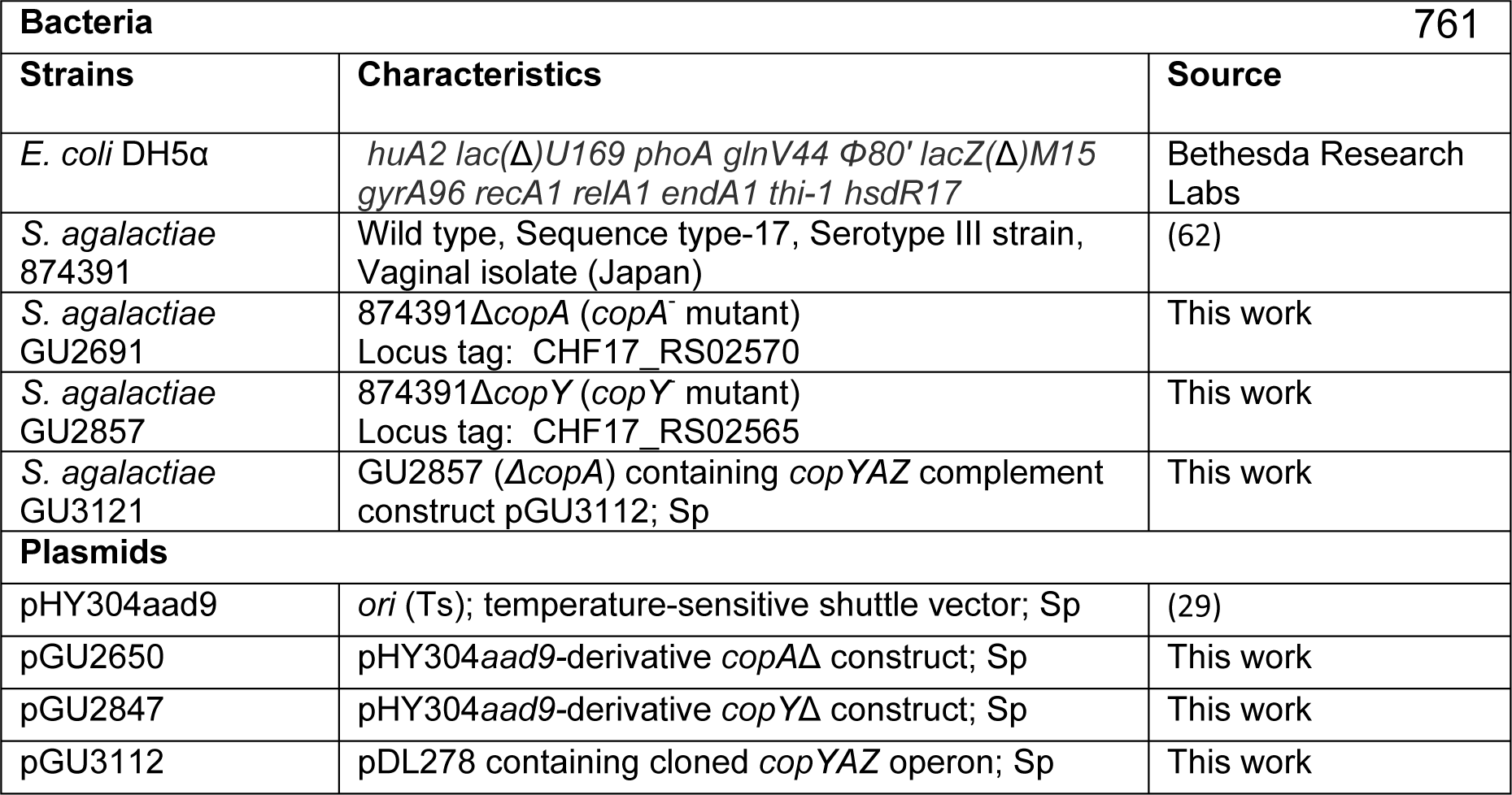
Bacterial strains and Plasmids.

### DNA extraction and genetic modification of GBS

Plasmid DNA was isolated using miniprep kits (QIAGEN), with modifications for GBS as described elsewhere (28). Deletions in *copA* (CHF17_00507 / CHF17_RS02570) and *copY* (CHF17_00506 / CHF17_RS02565) were constructed by markerless allelic exchange using pHY304aad9 as described previously (29). Plasmids and primers are listed in Table 2 and Table 3, respectively. Mutants were validated by PCR using primers external to the mutation site and DNA sequencing.

**TABLE 3.**
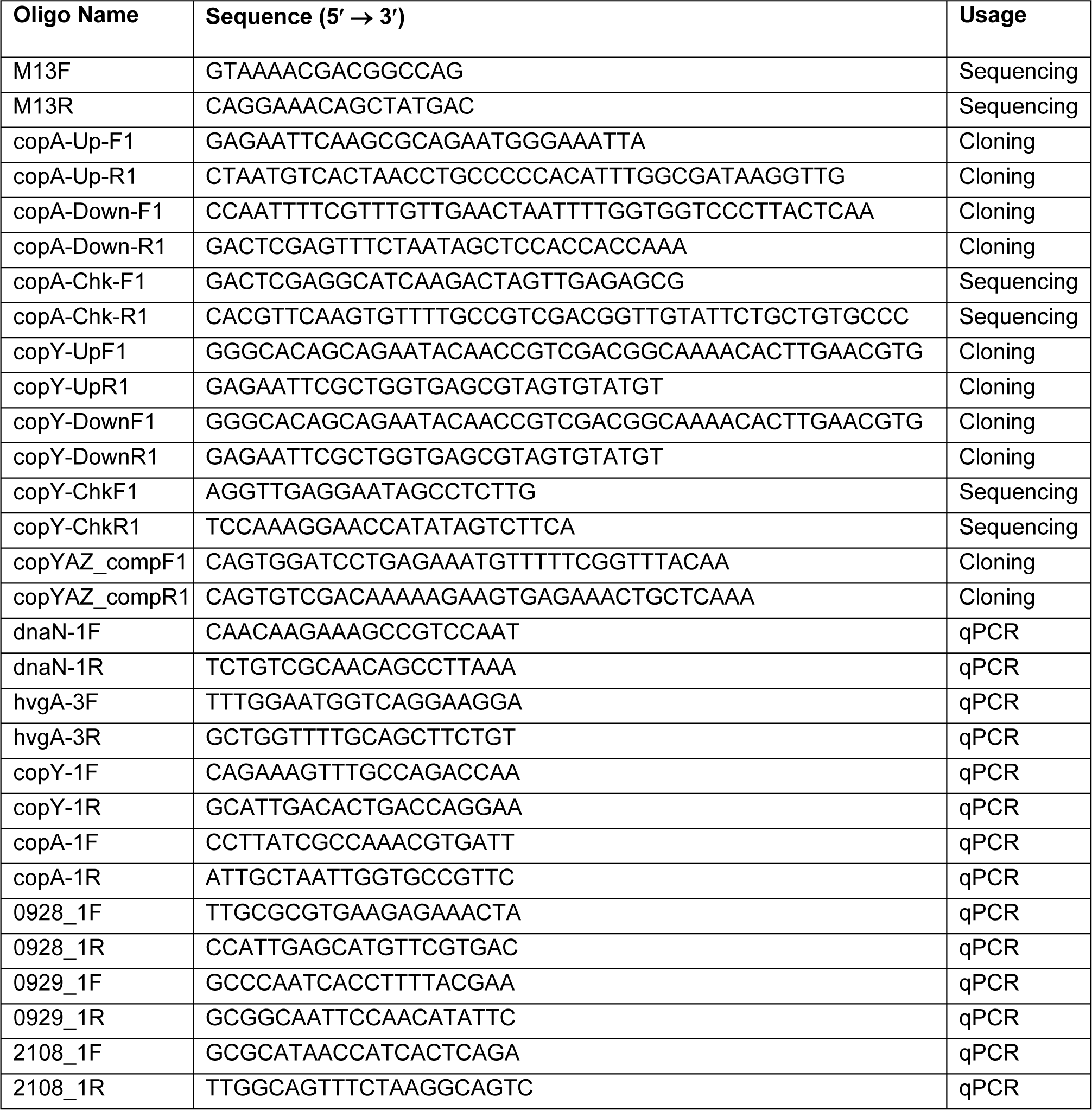
Primers.

### RNA extraction, qRTPCR

For Cu exposure experiments, 1mL of overnight THB cultures were back-diluted 1/100 in 100mL of THB (prewarmed at 37°C in 250mL Erlenmeyer flasks) supplemented with 0.25, 0.5, 1.0 or 1.5mM Cu. Cultures were grown shaking (200rpm) at 37°C; after exactly 2.5h, 10-50mL volumes containing approximately 500 million mid-log bacteria were harvested; RNA was preserved and isolated as described previously (30). RNA quality was analysed by RNA LabChip using GX Touch (Perkin Elmer). RNA (1000ng) was reverse-transcribed using Superscript IV according to manufacturer’s instructions (Life Technologies) and cDNA was diluted 1:50 in water prior to qPCR. Primers (Table 3) were designed using Primer3 Plus (31, 32) to quantify transcripts using Universal SYBR Green Supermix (Bio-Rad) using a Quantstudio 6 Flex (Applied Biosystems) system in accordance with MIQE guidelines (33). Standard curves were generated using five-point serial dilutions of genomic DNA (5-fold) from WT GBS 874391 (34). Expression ratios were calculated using C_T_ values and primer efficiencies as described elsewhere (35) using *dnaN*, encoding DNA polymerase III β-subunit as housekeeper.

### Whole bacterial cell metal content determination

Metal content in cells was determined as described (36) with minor modifications. Cultures were prepared essentially as described for *RNA extraction, qRTPCR* with the following modifications; THB medium was supplemented with 0.5 mM CuSO_4_ or not supplemented (Ctrl), and following exposure for 2.5h, bacteria were harvested by centrifugation at 4122 x g at 4°C. Cell pellets were washed 3 times in PBS + 5mM EDTA to remove extracellular metals, followed by 3 washes in PBS. Pelleted cells were dried overnight at 80°C and resuspended in 1mL of 32.5% nitric acid and incubated at 95°C for 1h. The metal ion containing supernatant was collected by centrifugation (14,000 x g, 30min) and diluted to a final concentration of 3.25% nitric acid for metal content determination using inductively coupled plasma optical emission spectroscopy (ICP-OES). ICP-OES was carried out on an Agilent 720 ICP-OES with axial torch, OneNeb concentric nebulizer and Agilent single pass glass cyclone spray chamber. The power was 1.4kW with 0.75L/min nebulizer gas, 15L/min plasma gas and 1.5L/min auxiliary gas flow. Cu was analysed at 324.75nm, Zn at 213.85nm, Fe at 259.94nm and Mn at 257.61nm with detection limits at <1.1ppm. The final quantity of each metal was normalised using dry weight biomass of the cell pellet prior to nitric acid digestion, expressed as µg.g^-1^dry weight. Baseline concentrations were determined to be 0.2 ± 0.08 µM Cu in THB medium, and 40 ± 4 nM Cu in CDM medium from at least three independent assays.

### Hydrogen Peroxide assay

Peroxide survival assays were based on prior studies (37, 38) with minor modifications to encompass using GBS cells that had been pre-grown in conditions of Cu stress. Overnight cultures of WT and mutants were grown in THB, cultures were back-diluted 1/100 into either fresh THB or THB supplemented with 0.5mM Cu and grown for exactly 2.5 h at 37°C with 200 rpm agitation. Bacteria were harvested by centrifugation at 4122 x g and washed twice in PBS and resuspended in assay buffer (0.1M sodium phosphate buffer pH 7.5). This was prepared by combining 41 mL of 0.2M dibasic sodium phosphate and 9 mL of 0.2M monobasic sodium phosphate in a volume of 200 mL.

GBS cells were diluted in assay buffer alone, or assay buffer containing 5 mM H_2_O_2_ (Sigma-Aldrich H1009), to a final concentration of ∼5x10^6^ CFU/mL. Cell suspensions +/-H_2_O_2_ were then incubated for 1h at 37°C and survival was monitored by serial dilution and plating for CFU/mL counts.

### RNA sequencing and bioinformatics

Cultures were prepared as described above for *RNA extraction, qRTPCR* to compare mid-log phase cells grown in THB + 0.5 mM Cu to THB without added Cu. RNase-free DNase-treated RNA that passed Bioanalyzer 2100 (Agilent) analysis was used for RNA sequencing (RNA-seq) using the Illumina NextSeq 500 platform. We used a Bacterial Ribosomal RNA (rRNA) Depletion kit (Invitrogen) prior to library construction, and TruSeq library generation kits (Illumina, San Diego, California). Library construction consisted of random fragmentation of the RNA, and cDNA production using random primers. The ends of the cDNA were repaired and A-tailed, and adaptors were ligated for indexing (with up to 12 different barcodes per lane) during the sequencing runs. The cDNA libraries were quantitated using qPCR in a Roche LightCycler 480 with the Kapa Biosystems kit (Kapa Biosystems, Woburn, Massachusetts) prior to cluster generation. Clusters were generated to yield approximately 725K–825K clusters/mm^2^. Cluster density and quality was determined during the run after the first base addition parameters were assessed. We ran single-end 75–bp sequencing runs to align the cDNA sequences to the reference genome. For data preprocessing and bioinformatics, STAR (version 2.7.3a) was used (parameters used: --outReadsUnmapped Fastx -- outSAMtype BAM SortedByCoordinate --outSAMattributes All) to align the raw RNA sequencing fastq reads to the WT *S. agalactiae* 874391 reference genome (34). HTSeq-count, version 0.11.1 (parameters used: -r pos -t exon -i gene_id -a 10 -s no -f bam), was used to estimate transcript abundances (39). DESeq2 was then used to normalized and test for differential expression and regulation following their vignette. Genes that met certain criteria (i.e. fold change of > ±2.0, q value (false discovery rate, FDR of <0.05) were accepted as significantly altered (40). Raw and processed data were deposited in Gene Expression Omnibus (accession no. GSE167895 for *S. agalactiae* 874391 Cu condition; GSE167894 (*S. agalactiae* 874391 control condition).

### Mammalian cell culture

U937 monocytes were differentiated into human monocyte-derived macrophages (MDMs) as follows. Briefly, monocytes were seeded (5x10^5^ per well such that 1x10^5^ adhere) into the wells of a 96-well tissue culture-treated plate (Falcon) essentially as described elsewhere (41, 42), except that U937 monocytes were differentiated by exposure to 30ng/mL phorbol 12-myristate 13-acetate (PMA) for 48h and cells subsequently rested in media without PMA for 72h to enhance morphological and phenotypic markers of MDMs (43). A multiplicity of infection (MOI) of 100 bacteria: macrophage for 1h was used in RPMI without antibiotics. Non-adherent bacteria were removed by five washes of 200µL PBS using a Well Wash Versa (Thermo Scientific). RPMI containing 250U/mL penicillin, streptomycin (Gibco) and 50µg/mL gentamicin (Sigma-Aldrich) were used for antibiotic protection assays to kill extracellular bacteria as described previously by incubating for 1h at 37°C in 5% CO_2_ (42). Samples were processed after 1h (time zero), 24h or 48h after infection, monolayers were washed five times with 200µL PBS and lysed by brief exposure to 50µL of 0.25% trypsin EDTA (Gibco) and 0.1% Triton-X-100 (10 min) prior to dilution with 150µL PBS and estimation of CFU/mL by serial dilution and plate counts on agar. Additional assays that ran in parallel were identical except that 20 µM Cu was added to RPMI culture media at all stages post PMA-treatment of U937 cells.

### Animals and Ethics statement

Virulence was tested using a mouse model of disseminated infection based on intravenous challenge with 10^7^ GBS (WT or ΔcopA) as described elsewhere (26). This study was carried out in accordance with the guidelines of the Australian National Health and Medical Research Council. The Griffith University Animal Ethics Committee reviewed and approved all experimental protocols for animal usage according to the guidelines of the National Health and Medical Research Council (approval: MSC/01/18/AEC).

### Statistical methods

All statistical analyses used GraphPad Prism V8 and are defined in respective Figure Legends. Statistical significance was accepted at P values of ≤0.05.

## Discussion

Transcriptional and cellular responses of bacterial pathogens to metal ions including Cu can influence host-pathogen interactions and thereby play a role in disease pathogenesis (4). The role of Cu homeostasis and detoxification in the biology of GBS have not hitherto been characterized, and no reports of a Cu stress response in this important human and animal pathogen are published. The principal finding of this study is the establishment of a transcriptional and cellular connection between the response to Cu stress in GBS and survival of the bacteria in conditions of Cu toxicity; this connection is mediated through *copA* and controlled through *copY,* and enables the bacteria to resist killing via Cu-mediated intoxication. Additionally, this study establishes that the connection between Cu stress responses in GBS and bacterial survival promotes virulence in the host during systemic, disseminated infection. The new insights into gene function in GBS viewed through the lens of the Cu stress transcriptome, combined with the findings of enhanced virulence elucidate molecular mechanisms that underpin GBS survival of intoxicating conditions, including those likely to be encountered in the host.

The transcriptional remodelling that occurs in GBS in response to Cu stress, as defined in this study on a global level, comprises an intriguingly constrained subset of genes. These findings indicate a tightly controlled system of transcriptional responses to Cu in GBS. Interestingly, equivalent low numbers of target genes were identified in previous transcriptional analyses of other streptococci exposed to Cu stress (14-16). Our findings are consistent with these prior reports, and support the notion that these transcriptional responses function in housekeeping or homeostasis to set a low limit of Cu availability in the cytoplasm (15). In GBS, *copA* is among the most strongly activated genes in the transcriptional response to Cu stress, and the mutational analysis performed in this study shows that *copA* is crucial for the bacteria to attain an essential Cu efflux response during Cu stress. However, *copA* is only one of an assembly of genes engaged by GBS during Cu stress and it is likely that other genes in the transcriptome contribute to Cu detoxification via additional means. For example, we detected up-regulation of putative metal transporters for Mn import (*mtsABC*, *mntH2*) along with concurrent down-regulation of a system that encodes Fe export machinery *(fetAB)*. These transcriptional insights are interesting because they hint at additional stress response mechanisms that occur during Cu stress in GBS, which extend beyond CopA and which need elucidation. Metal content analysis by ICP-OES showed that high Cu stress disrupts additional pools of intracellular trace metal, including for example causing modest elevations in levels of Fe along with major increases in Cu content. The consequences of these alterations in metal content in GBS for cell activity need further examination.

Transcription of metal ion-import and export genes, including those above-mentioned, is typically controlled by metal-dependent regulatory proteins termed metalloregulators that sense metal ion-bioavailability and work to maintain cellular metal homeostasis (44). In our study, we detected no major changes in the expression of genes predicted to encode metalloregulators, such as for transport of Mn (*mntR/mtsR*), Fe (*fur*), Zn (*adcR/sczA*) and for the sensing of peroxide (*perR*) in GBS undergoing Cu stress (25, 44). It would be of interest to elucidate whether such regulators undergo mis-metallation in GBS during Cu stress, which might occur due to excess Cu likely outcompeting Mn and Fe for binding sites in proteins; it is conceivable that Cu may displace metals from proteins as described in other studies (45); however, precisely if and how metalloregulators might respond to mis-metallation in GBS, if this occurs, will need to be elucidated in future.

In *S. pyogenes*, Mn uptake is facilitated by *mtsABC* and is protective against peroxide induced stress (25). Expression of *mtsABC* is up-regulated by Mn deficiency, and is controlled by the MtsR regulator (25, 46). Mn import and Fe efflux are co-ordinated in order to control the metalation of superoxide dismutase (25), which can use Mn or Fe at its catalytic site in streptococci; noting that Mn uptake can be disrupted by Zn (36). The consequences of Cu intoxication in relation to oxidative stress was explored using killing assays with hydrogen peroxide. These revealed reduced viability in *copA*-deficient GBS, but only in cells that had been pre-exposed to Cu. Taken with our observations that Cu accumulates significantly in this strain following Cu intoxication (and to a lesser extent Mn and Fe), these data hint at a role for oxidative damage as a killing mechanism. Fe can produce hydroxyl radicals and Cu(II) can be oxidised to Cu(I) in reactions with peroxide, both products of which are likely to damage bacterial cells. A system for the dual import of Mn and Fe in GBS is encoded by *mntH* that is regulated by pH (47). In some strains of GBS, including the hypervirulent ST17 lineage used in this study, two homologues of MntH exist, encoded by *mntH* and *mntH2*. This study demonstrates up-regulation of *mntH2*, but not *mntH*, in response to Cu, however the role of *mntH2* or its regulation are undefined in GBS. In *Bacillus subtilis* MntR co-ordinates Mn import (and efflux), including control of genes homologous to GBS *mntH* and *mntH2* that import Mn (48). This may provide context for elevated *mntH2* expression in the presence of modest Cu stress in our study, since Cu-bound MntR has a much lower affinity for binding its target operator sequence (14580210). We also note that *mtsABC* and *mntH2* are also up-regulated in concert with down-regulation of Fe-transporting *fetAB* in response to Zn stress in GBS (49). Together, our data indicate that modulation of *mtsABC*, *mntH2* and *fetAB* expression forms parts of a transcriptional signature of GBS exposed to Cu or Zn stress.

We observed no significant transcriptional response of *nrdD* expression in WT GBS exposed to Cu stress. In contrast, we detected a highly statistically significant, but modestly fold-change (up-regulated) response of the *nrdFIA* genes, encoding aerobic dNTP synthesis components (1.7-fold, FDR < 0.001; data not shown). These findings might hint at a role for Mn in the GBS Cu stress response given NrdD encodes an Fe-containing enzyme, whereas NrdF encodes a Mn-dependent pathway. The role of these genes in the GBS response to Cu stress requires further study. Modulation of putative Mn (*mtsABC*) and Fe (*fetAB*) transporters, as well as other proteins predicted to localise to the membrane (*pcl1*) or cell surface (*hvgA*) in GBS in response to Cu stress takes place in the absence of altered cellular Mn or Fe content in the WT strain. This leads us to suggest that examining the roles of targets identified in this study in the context of Cu transport is needed in concert with other metals (*e.g.,* Mn). Indeed, the ‘dearth of information’ on precisely how Cu is transported into bacterial cells from the external environment represents a central mystery for Cu trafficking in bacteria (50).

Our assays of bacterial growth *in vitro* in conditions of Cu stress demonstrate that GBS deficient in *copA* cannot grow as efficiently compared to the WT in conditions of elevated extracellular Cu. The differences noted in GBS growth between nutritionally rich (THB) and limited (CDM) media possibly reflect relative quantities of compounds that confer a protective advantage for survival during Cu stress, such as glutathione (15) or other thiol-containing amino acids that may interact with free Cu ions in solution. Glutathione is not included in CDM as a separate chemical constituent but the quantities of methionine, cystine and cysteine are 30, 62.6 and 50 mg.L^-1^. It is also possible that the levels of Cu utilized in our assays are distinct to those in host niches. Importantly, however, Cu exposure assays *in vitro* are almost certainly influenced by the compounds present in the medium that likely affect levels of Cu that become inhibitory (15). For example, buffering effects would likely occur in THB, which would affect the uptake of Cu by the bacteria. Our rationale for use of THB to analyse the impact of Cu toxicity on GBS is that this medium is a standard rich growth medium for GBS and is widely used for studies of this organism. The challenges inherent in such a rich media, including for example, buffering effects, led us to also examine the impact of Cu toxicity on GBS in defined CDM, which revealed, expectedly, a heightened susceptibility of the bacteria to the metal stress. Other similar studies have reported degrees of intoxication effects due to metal stress in different growth media, and our findings are consistent with these observations. Some of the Cu concentrations used in our study may be considered supra-physiological (1-1.5 mM) but the magnitude of these concentrations is likely much greater than the magnitude of ‘free’ Cu ions (not bound by other molecules) that would be present and available to react biologically. The concentration ranges used in the current study were informed by available literature along with measures of expression of *copA*, since we and others have shown that transcriptional activity of *copA* is influenced by cellular Cu. Thus, *copA* activity can be considered a *bona fide* reporter of free cellular Cu; or a surrogate marker for exposure to Cu stress. Consistent with this notion, we observed higher expression of *copA* (7.6 ± 2.3 fold up-regulated; not shown) in CDM with 100 μM than in THB with 500 μM Cu (4.9 ± 0.5 fold up; Figure 5). The Cu concentrations that are encountered by GBS at infection sites or in different intracellular compartments in host cells are unknown. Cu is elevated in the host at sites infected with *S. pyogenes* (15), including the blood and in infected skin lesions. For mycobacterial phagosomes, investigators have reported Cu concentrations of 25μM, and in *M. tuberculosis*-infected macrophages, intravacuolar concentrations of 0.4mM (51). Notwithstanding considerations of possible physiological ranges of Cu, the Cu exposure assays used in our study are beneficial to establish *bona fide* gene function and bacterial responses to a defined Cu stress condition.

Our analysis of Cu content in GBS cells exposed to Cu stress in defined conditions shows that GBS maintains low steady-state levels of cellular Cu, in the absence of excess extracellular Cu. In supplying extracellular Cu in excess, we show the level of cellular Cu content is increased. Our finding that *copY* functions to repress *copA* in the absence of Cu is consistent with previous reports in other bacteria (52, 53). Disrupting the genetic systems for Cu efflux in GBS, via mutation of the CopA exporter, or the CopY regulator, reveals divergent phenotypes that stem from loss of export or regulatory function, resulting in accumulation (Δ*copA*) or reduction (Δ*copY*) of cellular Cu. These phenotypes will be of interest to dissect in terms of the role of CopY in GBS biology in other models of infection and disease in the future. The results of the current study indicate that GBS might have a higher intrinsic level of resistance to Cu compared to other *Streptococcus* spp., including *S. pyogenes*. However, we did not directly compare different species and differences in experimental approaches or background solution, as our study demonstrates, might affect these assays. Thus, further work characterising relative resistance between streptococci more directly would be of interest to the field. The potential for Cu resistance mechanisms to confer heightened Cu tolerance in the context of the host environment is unexplored for GBS.

Studies have demonstrated increased Cu levels in the blood of humans during bacterial infection (54, 55), however most of the insight into Cu-driven antimicrobial responses is derived from studies of mammalian cells infected *in vitro*. In macrophages, bioavailability of Cu correlates with antibacterial activities (56), and Cu ‘hot spot’ formation mediates antimicrobial responses against intracellular bacteria (7). The cellular consequences of Cu stress to GBS, which likely encounters such stress in the host, remains undefined. Our findings based on *in vitro* infection macrophages showed no attenuation of GBS devoid of CopA in survival in host cells, which was surprising given the important role of Cu management in survival of other bacteria inside macrophages (6), and epithelial cells (12). Notably, however, intracellular survival of *Salmonella* deficient in *cueO*, which encodes an enzyme required for resistance to Cu ions, was not impaired in murine macrophages in a previous study (57), leading the authors to suggest multiple host factors are involved in clearance of the bacteria. In addition, *copA-*deficient *S. pyogenes* were not impaired for survival in assays with human neutrophils [15]. Our study is consistent with these findings. Other researchers have described the limitations of *in vitro* tissue culture monolayer assays for determining intracellular survival of bacteria in the context of Cu homeostasis (58).

Despite negative findings in macrophage monolayer assays *in vitro*, systemic infection of mice exposed a connection between the ability of GBS to generate a Cu management response via CopA and bacterial virulence *in vivo*. Here, *copA* was essential for GBS to fully colonize and survive in the blood, as well as in other tissues. In demonstrating a significant attenuation of GBS deficient in CopA to be fully virulent in mice, we suggest that Cu toxicity may represent a form of stress experienced by the bacteria *in vivo* during systemic infection. In other bacteria, including *Pseudomonas aeruginosa* and *Listeria monocytogenes*, compromised Cu transport leads to attenuation for colonization in various infection models (58, 59). Attenuation of GBS for colonization of the blood, liver and spleen indicates that Cu management in the bacterial cell is essential not only for efficient survival of the bacteria in the bloodstream but also for colonization of highly immunologically active tissues; *i.e.,* Kupffer cells and splenic lymphocytes for innate and adaptive immune responses, respectively. It is plausible that Cu stress that might be encountered by GBS *in vivo* could influence virulence factor function, such as the hypervirulence associated adhesin, HvgA, which we have shown is down-regulated under Cu stress. It would be of interest to analyse the effect of Cu transport deficiency in GBS in other relevant models of infection, including in vaginal colonization (60). Additional to defining the effects of metal homeostasis in GBS on the nature of infection and disease caused by this organism, small-molecules probes might hold promise for the identification of other molecular mechanisms of metal homeostasis in GBS, as reported for Gram-positive bacteria (61).

In summary, this study shows that management of Cu export in GBS is essential for the bacteria to survive in environments of Cu stress. Cu intoxication in GBS generates a transcriptional signature that includes activation of the *cop* operon to confer bacterial survival and virulence in stressful environments. The exact role for Cu ions as an antibacterial response against GBS warrants further investigation.

## Supporting information

Combined Supplementary Figures

## Acknowledgements

We thank Michael Crowley and David Crossman of the Heflin Centre for Genomic Science Core Laboratories, University of Alabama at Birmingham (Birmingham, AL) for RNA sequencing. We also thank Ryan Stewart at the School of Environment Analytical Chemistry Core Facility, Griffith University, for ICP-OES. This work was supported by a Project Grant from the National Health and Medical Research Council (NHMRC) Australia (APP1146820 to GCU).

