## Supplementary material for "Copper intoxication in group B *Streptococcus* triggers transcriptional activation of the *cop* operon that contributes to enhanced virulence during acute infection": Combined Supplementary Figures

**Supplementary Figure Legends**.

**Supplementary FIG S1.** Final biomass yield comparisons of WT, Δ*copA* and Δ*copY* strains following growth for 18h in THB supplemented with 1.5 mM Cu. Bars show mean attenuance (*D*, 600nm) and S.E.M of 3 independent experiments. The groups were compared by One-way ANOVA with Holm-Sidak multiple comparisons (*** P <0.001).

**Supplementary FIG S2.** Growth curve analyses of GBS cultured in CDM (A), and CDM supplemented with 0.2 mM Cu (B), 0.5 mM Cu (C) or 1.0 mM Cu (D), comparing WT and Δ*copA* strains (and complemented strain Δ*copA::copYAZ*). Points and bars show mean and S.E.M of several independent experiments (5 for WT and *copA*, 2 for complemented strain) monitoring attenuance at 600nm.

**Supplementary FIG S3.** Cellular metal content during high Cu stress.  Intracellular accumulation of Cu, Fe, Mn and Zn was compared with and without Cu supplementation (1.5 mM) in WT GBS. Bars show mean S.E.M of four independent experiments. The groups were compared by One-way ANOVA with Holm-Sidak multiple comparisons (* P < 0.05, ** P < 0.01, *** P <0.001).

**Supplementary FIG S4.** Growth in the presence of Cu enhances susceptibility of Δ*copA* GBS to $H_2O_2$. WT, Δ*copA* and Δ*copA::copA* (Δ*copA*+C) strains were grown to mid-log phase in THB or THB supplemented with 0.5 mM Cu and then subjected to $H_2O_2$ for 1h. Control incubations in buffer without $H_2O_2$ for all strains contained $4.2 \pm 0.4 \times 10^7$ CFU/mL. The groups were compared by One-way ANOVA with Holm-Sidak multiple comparisons (** $P < 0.01$).

**Supplementary FIG S5.** Recovery of WT (grey circles) and Δ*copA* (blue diamonds) GBS (orange triangles) in a mouse model of disseminated infection. C57BL/6 mice (6-8 weeks old) were intravenously injected with $10^7$ bacteria and CFU were enumerated in heart, lungs, bladder, brain and kidneys at 24h post infection. Counts were normalized using tissue mass in g. Viable Cell counts of 0 CFU/mL were assigned a value of 1 to enable visualisation on $\log_{10}$ y-axes. Lines and bars show median and interquartile ranges. Data are pooled from 2 independent experiments each containing n=10 mice.

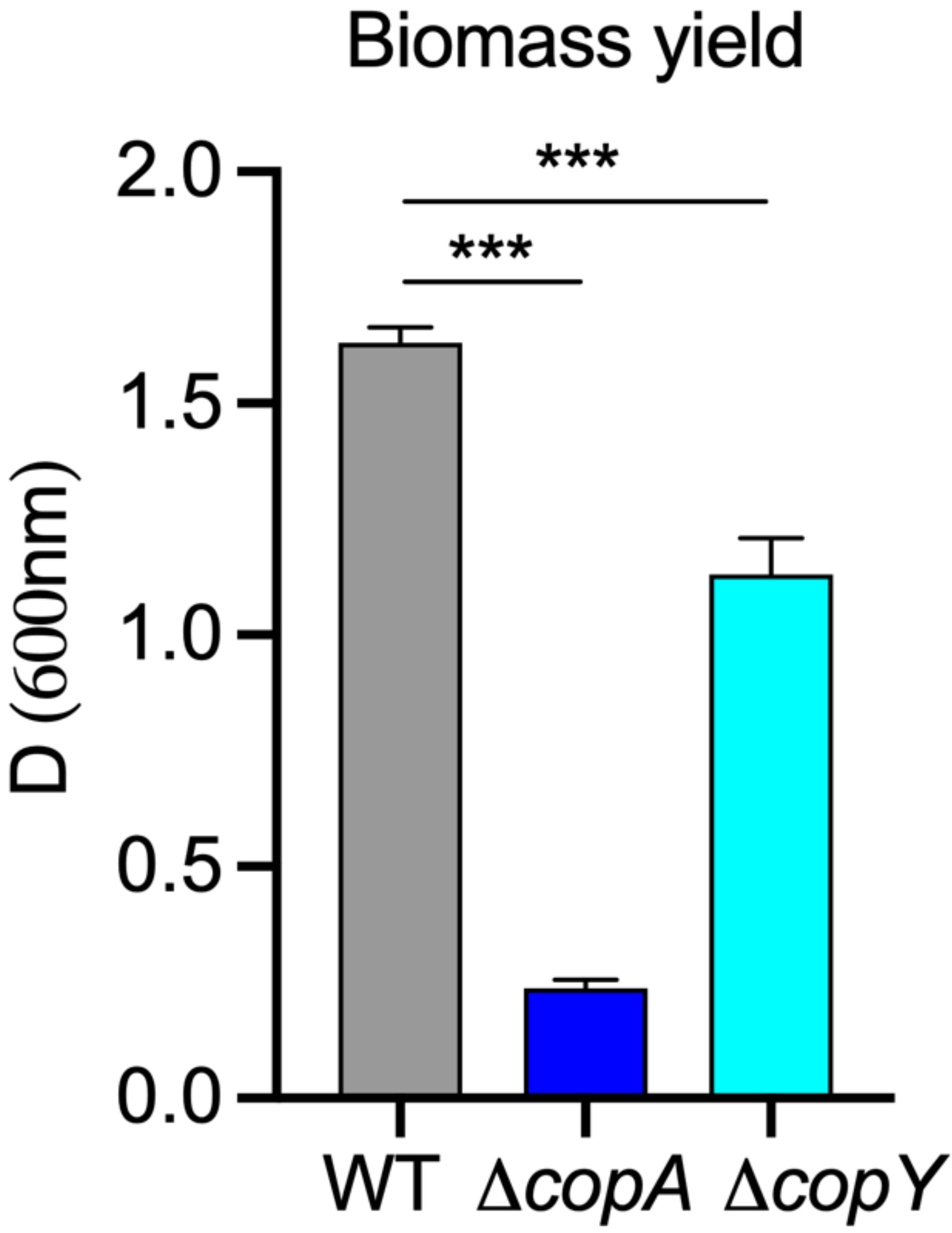

Biomass yield
D (600nm)
2.0
1.5
1.0
0.5
0.0
***
***
WT
ΔcopA
ΔcopY

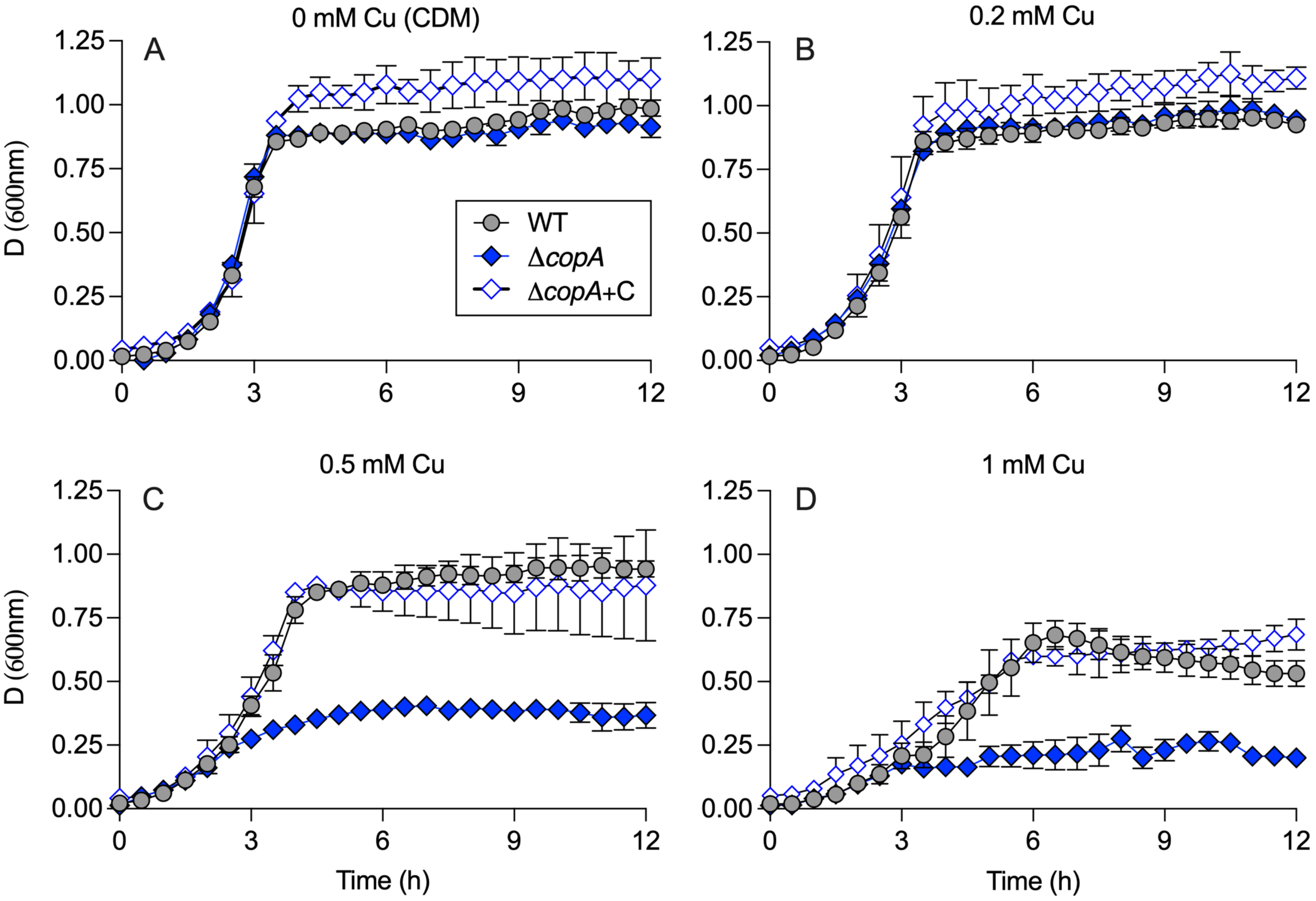

0 mM Cu (CDM)
A
0.2 mM Cu
B
0.5 mM Cu
C
1 mM Cu
D
WT
ΔcopA
ΔcopA+C
D (600nm)
Time (h)
0.00
0.25
0.50
0.75
1.00
1.25
0
3
6
9
12

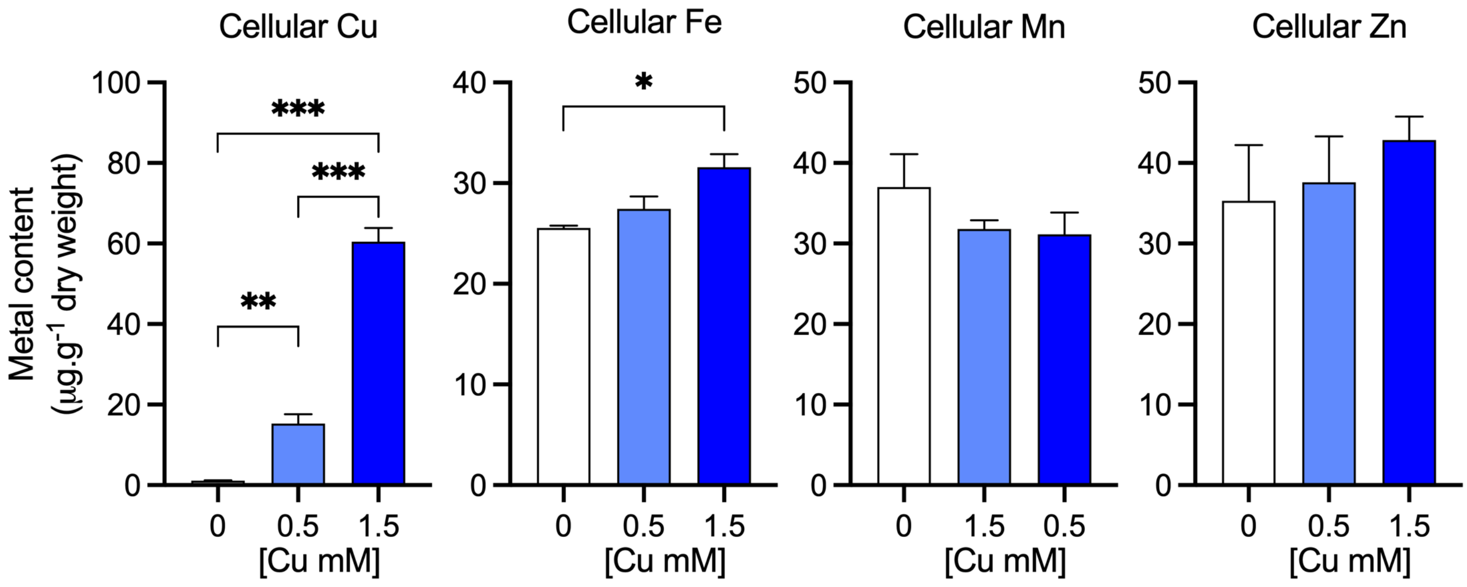
Cellular Cu
Cellular Fe
Cellular Mn
Cellular Zn
Metal content
(µg.g-1 dry weight)
***
***
**
*
[Cu mM]
0
0.5
1.5

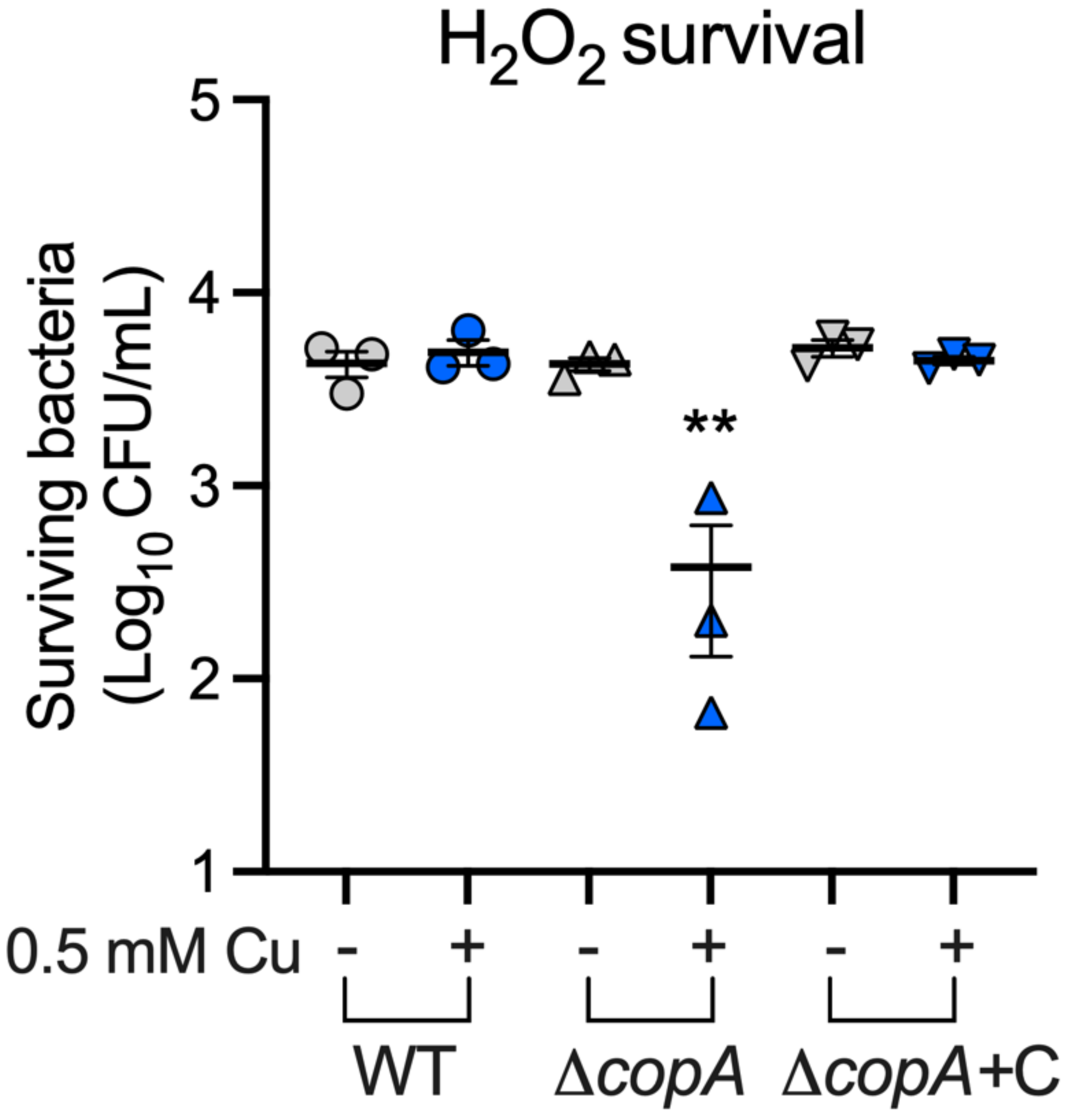

H2O2 survival
Surviving bacteria
(Log10 CFU/mL)
5
4
3
2
1
**
0.5 mM Cu
-
+
-
+
-
+
WT
ΔcopA
ΔcopA+C

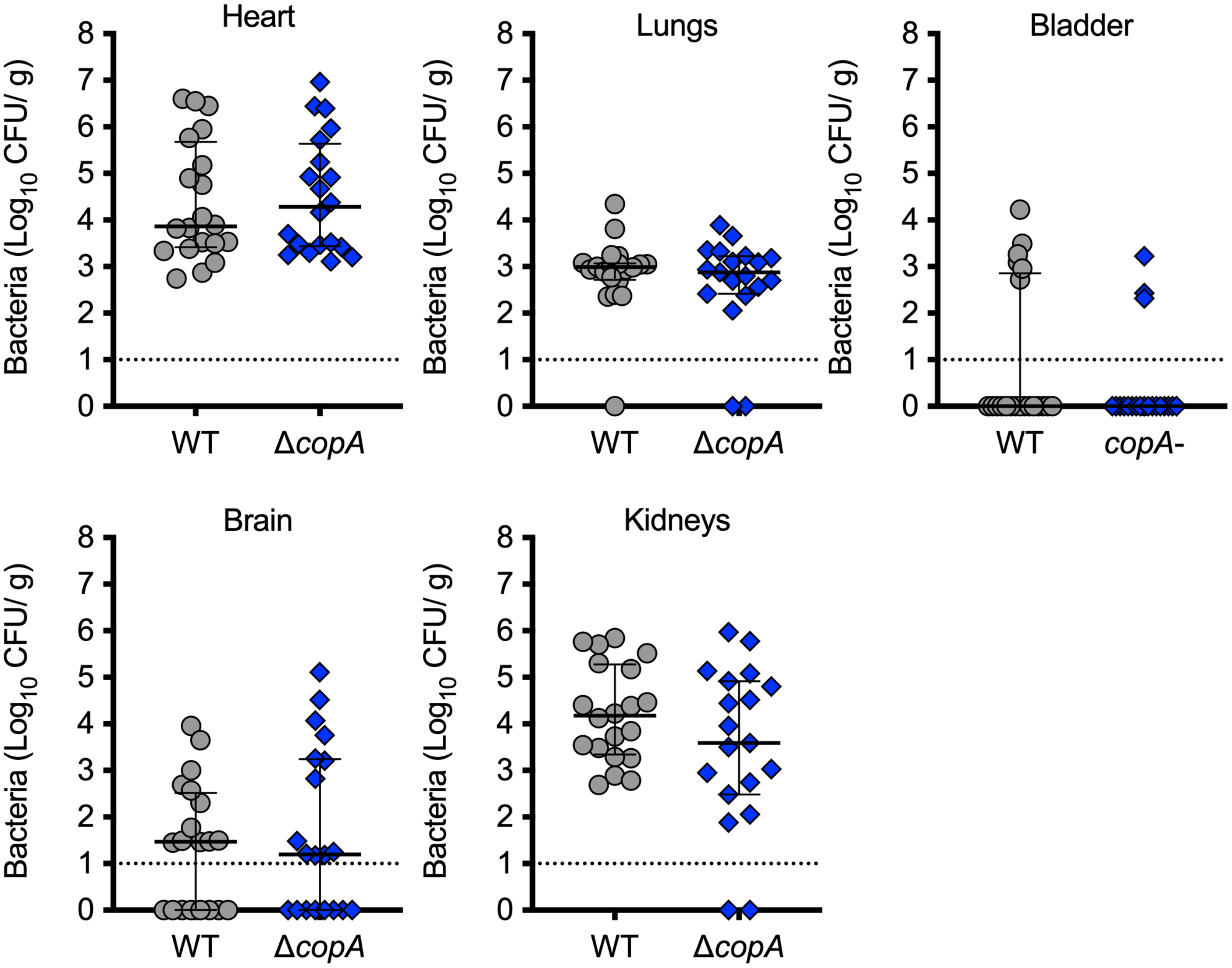
Heart
Lungs
Bladder
Brain
Kidneys
Bacteria (Log$_{10}$ CFU/ g)
8
7
6
5
4
3
2
1
0
WT
ΔcopA
copA-
